# Plzf mediates a switch between Fgf signalling regimes in the developing hindbrain

**DOI:** 10.1101/2022.09.23.509139

**Authors:** Sami A. Leino, Sean C. J. Constable, Andrea Streit, David G. Wilkinson

## Abstract

Developing tissues are sequentially patterned by extracellular signals that are turned on and off at specific times. In the zebrafish hindbrain, fibroblast growth factor (Fgf) signalling has different roles at different developmental stages: in the early hindbrain, transient Fgf3 and Fgf8 signalling from rhombomere 4 is required for correct segmentation, whereas later, neuronal Fgf20 expression confines neurogenesis to specific spatial domains within each rhombomere. How the switch between these two signalling regimes is coordinated is not known. We present evidence that the promyelocytic leukaemia zinc finger (Plzf) transcription factor is required for this transition to happen in an orderly fashion. Plzf expression is high in the early anterior hindbrain, then gradually upregulated posteriorly and confined to neural progenitors. In mutants lacking functional Plzf, *fgf3* expression fails to be downregulated and persists until a late stage, resulting in excess and more widespread Fgf signalling during neurogenesis. Accordingly, the spatial pattern of neurogenesis is disrupted in *plzf* mutants. Our results reveal how the distinct stage-specific roles of Fgf signalling are coordinated in the zebrafish hindbrain.

## Introduction

The hindbrain is an embryonic structure that gives rise to the cerebellum, pons and medulla of the vertebrate central nervous system (CNS). It is transiently subdivided into seven or eight segments called rhombomeres along its antero-posterior axis; this segmental organisation underlies the generation of the various neuronal subtypes that arise from the hindbrain (Lumsden and Keynes, 1989; Marin and Puelles, 1995). Each rhombomere acquires a distinct identity on the basis of intercellular signalling and gene regulatory interactions (reviewed in Frank and Sela-Donenfeld, 2019; Krumlauf and Wilkinson, 2021), which in turn forms the basis for the identity of hindbrain-derived neurons (reviewed in Philippidou and Dasen, 2013). After the demarcation of segmental identities, rhombomere boundaries (the interfaces between two adjacent rhombomeres) acquire properties distinct from rhombomere centres (Lumsden and Keynes, 1989; Trevarrow et al., 1990; Guthrie and Lumsden, 1991; Heyman et al., 1995). In zebrafish, hindbrain boundaries are induced at segmental interfaces by a process involving mechanical tension (Cayuso et al., 2019) and in turn regulate the spatial pattern of neuronal differentiation by local inhibition of neurogenesis (Cheng et al., 2004; Voltes et al., 2019) and by signalling to non-boundary populations (Riley et al., 2004; Gonzalez-Quevedo et al., 2010; Terriente et al., 2012).

Fgf signalling has different functions in the hindbrain depending on the developmental stage. In zebrafish, rhombomere 4 (r4) acts as a signalling centre during early hindbrain development: *fgf8*, and later *fgf3*, are expressed in r4 from late gastrulation until mid-somitogenesis and regulate gene expression in r3 and r5, pointing to a role for Fgf signalling in regulating rhombomere identity (Maves et al., 2002; Walshe et al., 2002; Wiellette and Sive, 2004). At later stages, after downregulation of segmental *fgf8* and *fgf3* expression, Fgf20 signalling from neurons located in rhombomere centres antagonises neurogenesis and may have a role in promoting gliogenesis (Gonzalez-Quevedo et al., 2010; Esain et al., 2010; Tambalo et al., 2020). Since neuronal differentiation is also inhibited at rhombomere boundaries (Cheng et al., 2004; Voltes et al., 2019), neurogenesis only takes place in stripes between rhombomere centres and boundaries. Thus, in the zebrafish hindbrain, early Fgf signalling regulates rhombomere identity, whereas late Fgf signalling establishes the spatial pattern of neuronal differentiation, and these two functions involve different ligands this change is reflected in the expression pattern of the Fgf target gene *etv5b*, which is expressed first in a rhombomere-specific pattern, and later confined to segment centres (Münchberg et al., 1999; Roehl and Nüsslein-Volhard, 2001; Esain et al., 2010; Gonzalez-Quevedo et al., 2010).

In amniotes, the early patterning of rhombomeres is also regulated by Fgf signalling. In chick, the expression of r3-r6 markers is dependent on Fgf receptor signalling (Marin and Charnay, 2000; Aragon et al., 2005; Aragon and Pujades, 2009) and the Fgf3 ligand specifically (Weisinger et al., 2008; Weisinger et al., 2010). Fgf3 itself is initially expressed in a dynamic, segment-specific pattern and is later restricted to rhombomere boundaries in chick (Mahmood et al., 1995; Weisinger et al. 2008, 2010, 2012; Sela-Donenfeld et al., 2009) and mouse (Wilkinson et al., 1988; Mahmood et al., 1996). In the chick hindbrain, rhombomere boundaries act as pools of neural progenitors (Peretz et al., 2016) and the expression of neuronal markers at the boundaries is dependent on Fgf3 function (Weisinger et al., 2012).

These observations suggest that, in both fish and amniotes, there is a switch during hindbrain development from an early pattern of Fgf signalling conferring segmental identity to a late regime of Fgf expression regulating neurogenesis in specific, sub-rhombomeric domains. Nonetheless, the differences in the ligands involved and their expression patterns point to species-specific features in the regulation of Fgf signalling in the hindbrain. In chicken, BMP signalling (Weisinger et al., 2008) as well as an unidentified signal from the boundaries (Sela-Donenfeld et al., 2009) have been implicated in the downregulation of *Fgf3* outside the boundaries; however, how the switch between two different patterns of Fgf signalling is coordinated is not understood.

The promyelocytic leukaemia zinc finger (Plzf) protein regulates a number of different processes during normal development. Plzf opposes differentiation and maintains a progenitor state in haematopoietic and spermatogenetic stem cells (Shaknovich et al., 1998; Buaas et al., 2004; Costoya et al., 2004) but promotes differentiation of megakaryocytes (Labbaye et al., 2002), chondrogenesis (Liu et al., 2011) and osteogenesis (Agrawal Singh et al., 2019). Plzf is expressed widely in the embryonic amniote CNS, including the hindbrain (Avantaggiato et al., 1995; Cook et al., 1995; Tailor et al., 2013), as well as in the early neural epithelium in zebrafish (Sobieszczuk *et al*., 2010). In addition, Plzf is expressed specifically in rosette-forming human embryonic stem cell-derived neural stem cells, a stage exhibiting a wide differentiation potential (Elkabetz et al., 2008). In mouse and chicken embryos, *PLZF* mRNA is initially enriched in even-numbered rhombomeres and is gradually confined to rhombomere boundaries as development proceeds (Cook et al., 1995), suggesting potential roles in both hindbrain patterning and regulation of neural differentiation. However, there is no evidence from functional studies for regulation of hindbrain development by Plzf. In zebrafish, Plzf acts as an antagonist of neural differentiation during the development of primary neurons, and its degradation is required for neurogenesis to proceed (Sobieszczuk et al., 2010). In chicken and mouse, Plzf maintains spinal cord neural progenitors in an undifferentiated state by positively regulating Fgf receptor expression (Gaber et al., 2013). These results imply a conserved role for Plzf in neural progenitor maintenance.

Here, we examine the expression and function of Plzf in the zebrafish embryonic hindbrain. We find that Plzf has a dynamic and rhombomere-specific pattern of expression and that Plzf expression is largely confined to neural progenitors and absent from post-mitotic neurons. To assess the function of Plzf in hindbrain development, we generated Plzf double mutants, which reveal a role for Plzf in downregulating the early segment-specific expression of *fgf3*, required for the correct organisation of neurogenesis at later stages. Our results point to a previously undescribed role for Plzf in regulating Fgf ligand expression and show that Plzf function is essential for coordinating the switch between early and late patterns of Fgf signalling in the zebrafish hindbrain, which may shed light on the differences in how this process is regulated in fish and amniotes.

## Results

### Dynamic expression of Plzf in the zebrafish hindbrain

Previous studies have shown that the two *plzf* paralogues, *plzfa* and *plzfb*, are expressed in the zebrafish CNS, including a part of the hindbrain, at 10-12 hpf, but have not examined their temporal and spatial pattern in detail, particularly at later stages. To address this, we used hybridization chain reaction (HCR; Choi et al., 2018) in situ hybridization (ISH) to detect *plzfa* and *plzfb*, between 14 hpf, (10-somite stage) and 24 hpf, by which stage neurogenic domains are being established in the hindbrain (Gonzalez-Quevedo et al., 2010). To ascertain the identity of individual rhombomeres, we used a transgenic reporter line that expresses Citrine in rhombomeres 3 and 5 (r3 and r5; Addison et al., 2018). At 14 hpf and 16 hpf, *plzfa* is expressed widely in the hindbrain, with somewhat lower levels in r7 (Fig. 1 A, D). *plzfb* has a more rhombomere-specific expression pattern at 14 hpf, with higher levels of expression in r1 and r5-r7 as well as the anterior spinal cord (Fig. 1B). At 16 hpf, *plzfb* levels are higher in r2-r4, leading to a more uniform pattern of expression in the hindbrain (Fig. 1E).

**Figure 1.**
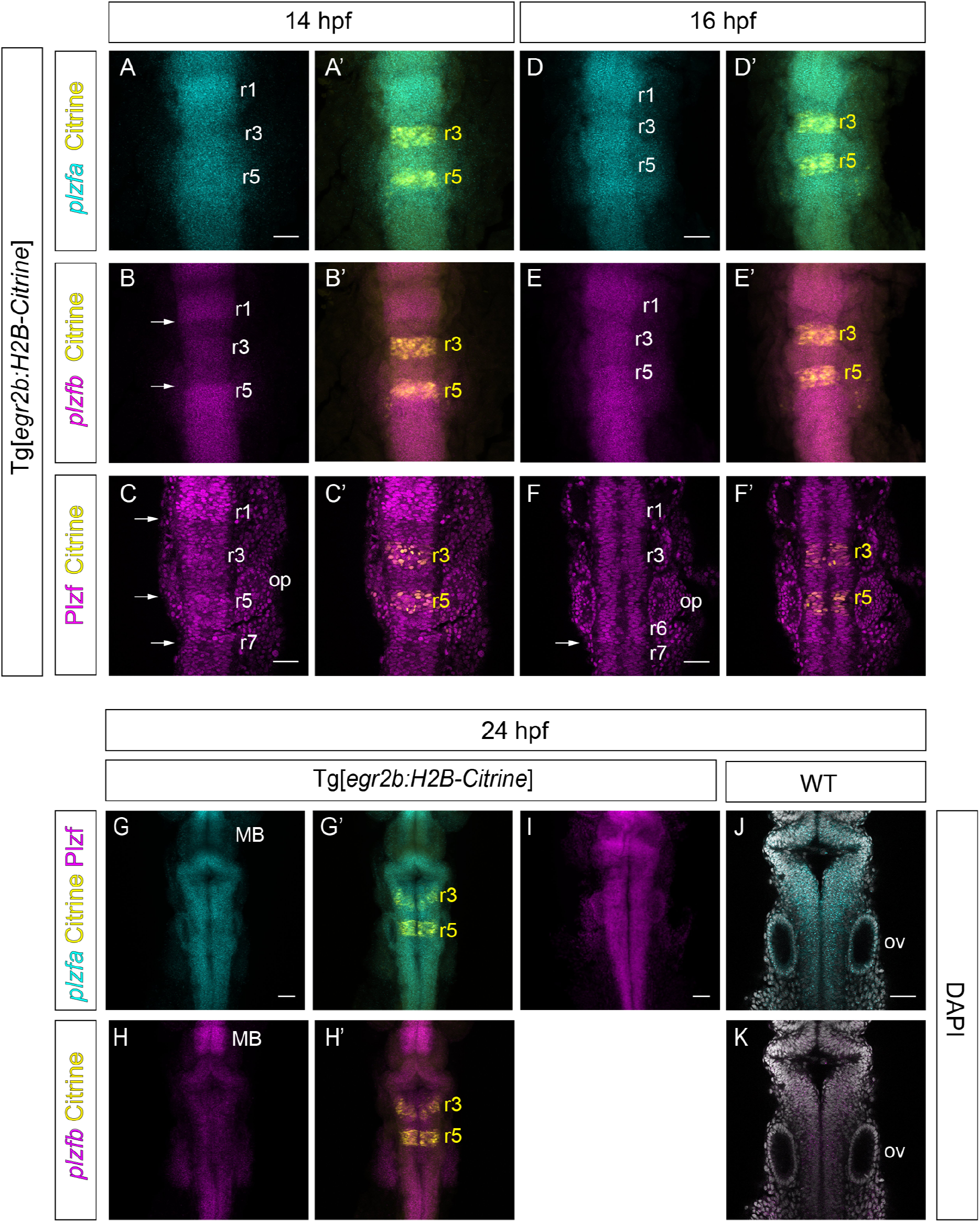
Expression of *plzfa/b* mRNA and Plzf protein in the zebrafish hindbrain. (**A,B; D,E; G,H**) HCR for *plzfa* and *plzfb* at 14, 16, and 24 hpf. Sum projections through the Z-stack. The Tg[*egr2b:H2B-Citrine*] line expresses Citrine in rhombomeres 3 and 5. Arrows indicate boundaries between high and low levels of *plzfb* expression. (**C,F,I**) Immunofluorescent staining with anti-Plzf antibody (see materials and methods) at 14, 16 and 24 hpf. (C,F) slices from confocal Z-stack; (I): Sum projection through Z-stack. Arrows indicate boundaries between high and low levels of Plzf expression. (**J,K**) HCR for *plzfa* and *plzfb* in a 24 hpf wild-type embryo. Slices from confocal Z-stack. Abbreviations: r, rhombomere; MB, midbrain; ov, otic vesicle. Scale bars: 50 µm

At 24 hpf, *plzfa* is expressed widely across the hindbrain and the anterior spinal cord, whereas *plzfb* levels appear low in the hindbrain (Fig. 1G, H). Additionally, *plzfb* mRNA is enriched in the midbrain in a domain with low levels of *plzfa* expression (Fig. 1G, H). At this stage, *plzfa* is also expressed in the otic vesicle, while *plzfb* levels are very low in this tissue (Fig. 1J, K.)

To complement the mRNA expression patterns, we stained for Plzf protein expression using an antibody that recognizes both Plzfa and Plzfb proteins (Fig. S1). Immunofluorescence reveals that, at 14hpf, Plzf protein is expressed at high levels in r1 and the anterior CNS as well as r5-r6 (Fig. 1C). By 16 hpf, Plzf protein levels are more uniform in the anterior hindbrain, whereas r7 and the anterior spinal cord show relatively lower levels of expression (Fig. 1F). This pattern of expression may be accounted for by higher Plzf protein levels in segments in which both *plzfa* and *plzfb* mRNAs are expressed. Finally, by 24 hpf, Plzf is present at uniform levels across the whole hindbrain and anterior spinal cord (Fig. 1I). Together, these results reveal that the Plzf expression begins in the anterior hindbrain and later expands into the posterior hindbrain and anterior spinal cord.

### Expression of Plzf in relation to neurogenesis

Since Plzf has been implicated in progenitor maintenance and repressing neuronal differentiation, we characterised its expression in relation to neurogenesis. We combined Plzf immunostaining with co-staining for neuronal and neural progenitor markers at 24 and 44 hpf, stages which fall within the major phase of neurogenesis in the zebrafish hindbrain (Lyons et al., 2003). At 24 hpf, the ventricular zone, which contains the neural progenitors, occupies most of the hindbrain. Plzf protein is present throughout the hindbrain ventricular zone but absent from the HuC/D-positive post-mitotic neurons (Fig. 2A-F). At 44 hpf, the mantle zone containing post-mitotic neurons has grown at the expense of the ventricular zone, and the latter has acquired a characteristic ‘T-shape’ (Lyons et al., 2003). Plzf is expressed throughout the ventricular zone but is absent from the mantle zone, except for a few post-mitotic neurons (Fig. 2J-L).

**Figure 2.**
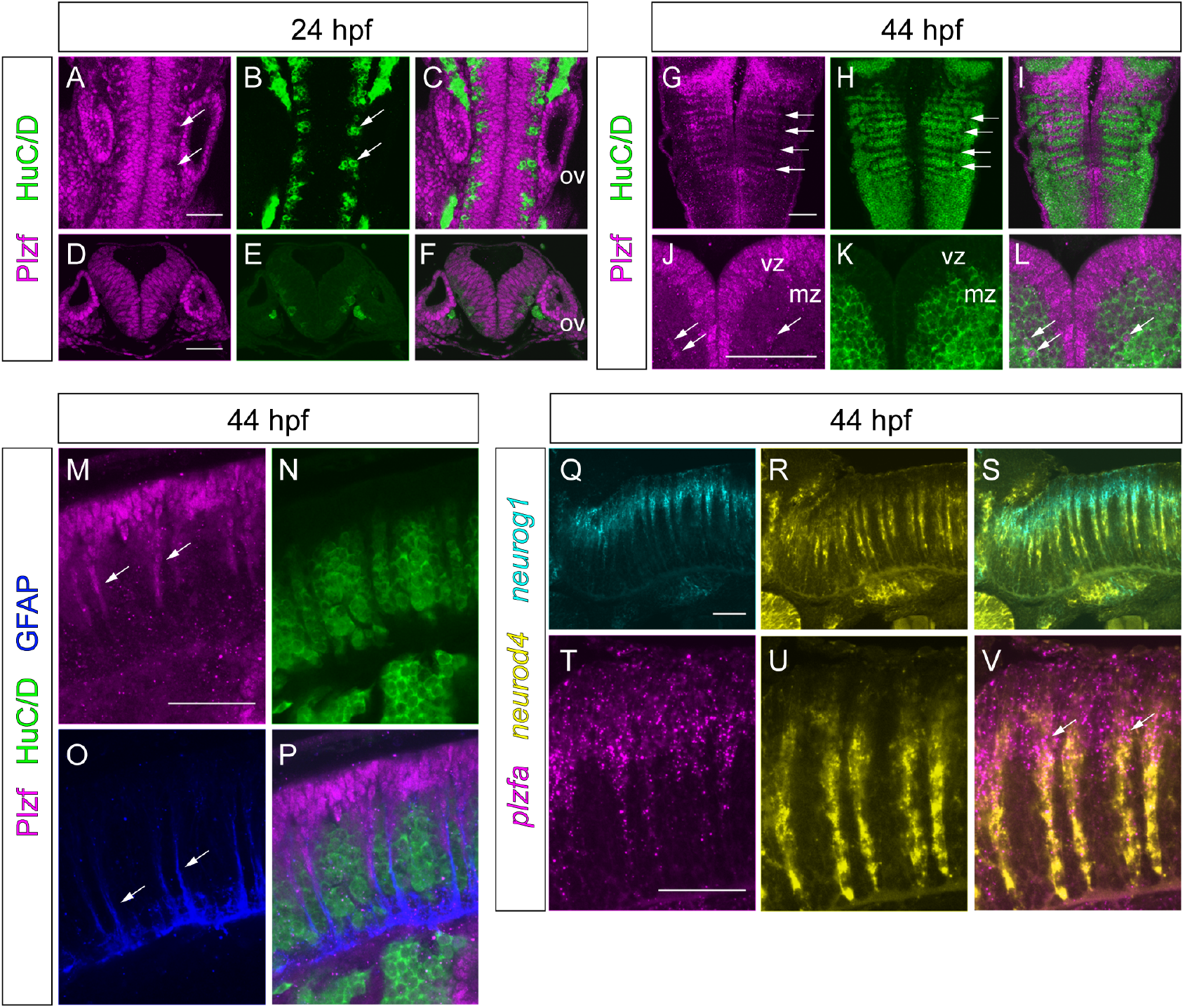
Expression of Plzf during neurogenesis. (**A**-**L**) Immunofluorescence for Plzf and HuC/D at 24 and 44 hpf. (**A**-**C**; **G**-**I**) dorsal views and (**D**-**F**; **J**-**L**) transverse sections of the hindbrain. Arrows indicate post-mitotic neurons (**A, B**), neurogenic zones (**G, H**) or Plzf + HuC/D double positive cells (**J,L**). (**M**-**P**) Immunofluorescence for Plzf, HuC/D and GFAP at 44 hpf, side views of the hindbrain, anterior to the left. Arrows indicate Plzf-positive migrating progenitors (**M**) or GFAP-positive glial fibres (**O**). (**Q**-**V**) Two-colour fluorescent ISH for *neurog1* and *neurod4* (**Q-S**) or *plzfa* and *neurod4* (**T-V**) at 44 hpf. Side views of the hindbrain, anterior to the left. Arrows indicate overlap of *plzfa* and *neurod4* expression. All images are slices from confocal Z-stacks. Abbreviations: ov, otic vesicle; mz, mantle zone; vz, ventricular zone. Scale bars: 50 µm

At the 44 hpf stage, neurogenesis in the hindbrain is localized to narrow stripes between the rhombomere centres and boundaries we term neurogenic zones (Gonzalez-Quevedo et al., 2010). The zones between segment centres and boundaries contain radial glial fibres that extend from the ventricular zone to the pial surface of the mantle zone (Trevarrow et al., 1990) and and create gaps in HuC/D staining (Fig. 2H). Cells in these gaps express Plzf (Fig. 2G-I). Side views of the hindbrain show Plzf staining in stripes of cells that extend ventrally from the ventricular zone and are negative for HuC/D staining (Fig. 2M, N). Co-staining for glial fibrillary acid protein (GFAP) reveals that the Plzf-expressing cell stripes coincide with glial fibres (Fig. 2M-P). Furthermore, most hindbrain neuronal progenitors at 44 hpf are radial glia (Lyons et al., 2003). Our observations therefore suggest that Plzf is expressed in neuronal progenitors in the ventricular zone and at early stages of migration from the ventricular zone to the mantle zone as they differentiate into neurons.

To pinpoint the state of differentiation of Plzf-expressing cells further, we examined the expression of *plzfa* mRNA in relation to proneural gene expression. *neurog1* is expressed in undifferentiated progenitors and cells that have started to differentiate, whereas the intermediate proneural gene *neurod4* is expressed later in the neurogenic cascade in cells that have migrated out of the ventricular zone and occupy a more basal position (Fig. 2Q-S). *plzfa* mRNA is detected in the ventricular zone as well as in early migrating progenitors, where its expression is apical to that of *neurod4* with a region of overlap (Fig. 2T-V), indicating co-expression of *plzfa* and *neurog1*. Taken together, these results show that Plzf is expressed in both undifferentiated and early differentiating neuronal progenitors, downregulated upon migration out of the ventricular zone, and mostly absent from post-mitotic neurons.

### The patterning of neurogenic zones is disrupted in Plzf mutants

We assessed the role of Plzf in hindbrain neurogenesis by examining proneural gene expression at 30 hpf in embryos lacking Plzf. To this end, we took advantage of transcription activator-like effector nuclease (TALEN) -mediated targeted mutagenesis to generate zebrafish mutant for both *plzf* paralogues (*plzfa*^*-/-*^;*plzfb*^*-/-*^). The mutations are predicted to generate null alleles, and indeed, no Plzf protein could be detected in double homozygous embryos after staining with the Plzf antibody (data not shown). At this stage, the neurogenic zones of the hindbrain are well delineated and can be visualised by detection of *neurod4* mRNA, whereas rhombomere centres are marked by expression of the ETS transcription factor *etv5b*, a target of Fgf signalling (Münchberg et al., 1999; Raible and Brand, 2001; Roehl and Nüsslein-Volhard, 2001). Co-staining of *plzfa*^*-/-*^;*plzfb*^*-/-*^ and sibling control embryos for *etv5b* and *neurod4* mRNA revealed ectopic *etv5b* expression in double mutants (Fig. 3A, E). In the sibling control, *etv5b* and *neurod4* are mostly complementary in the hindbrain (Fig. 3A-D), whereas both transcripts overlap extensively in the Plzf double mutant hindbrain and are expressed more uniformly instead of the strongly patterned expression seen in the sibling controls (Fig. 3E-H).

**Figure 3.**
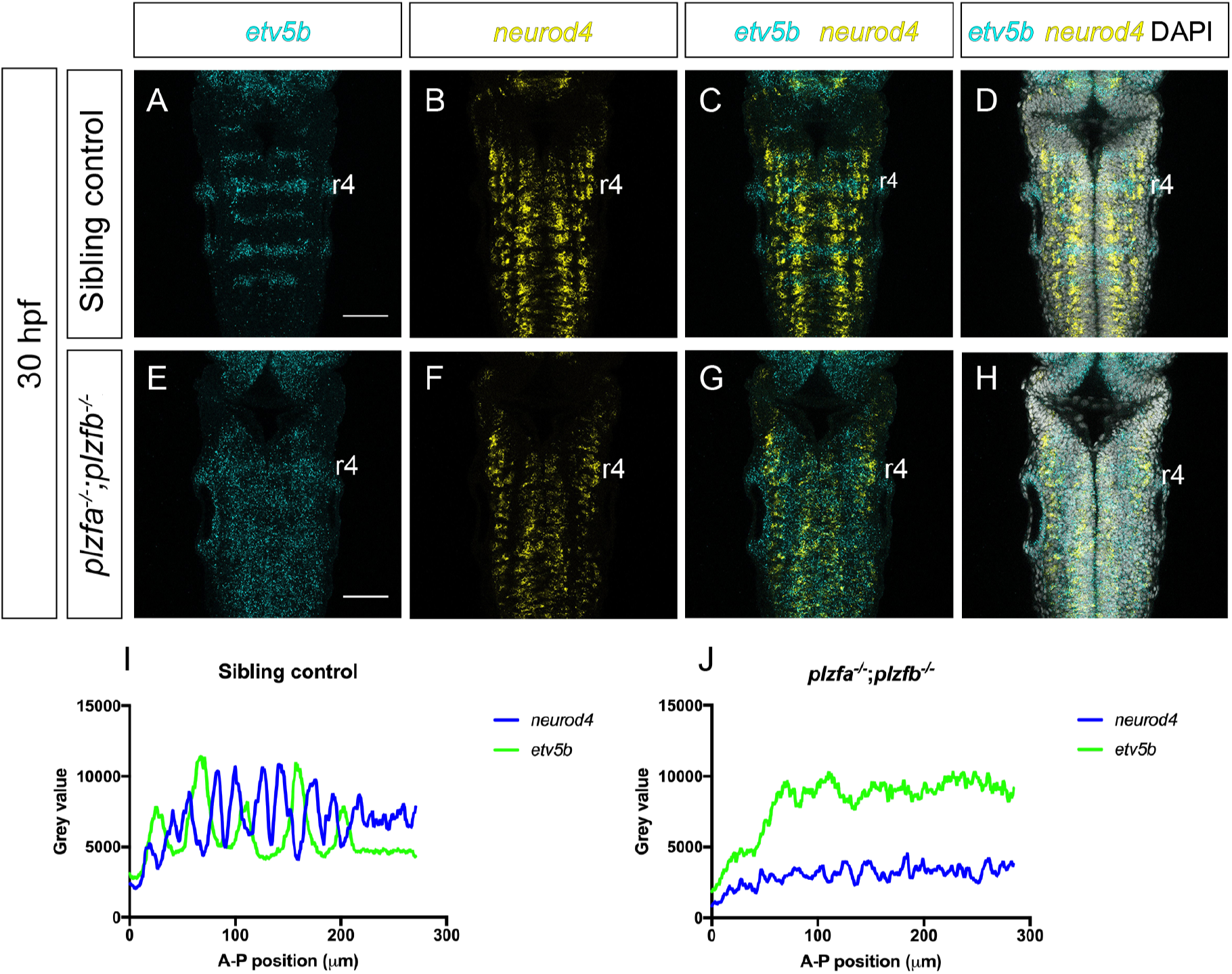
Spatial pattern of neurogenesis in *plzf* mutants. (**A-H**) HCR for *etv5b* and *neurod4* in 30 hpf sibling control (**A**-**D**; n=16) and *plzfa*^*-/-*^;*plzfb*^*-/-*^ (**E**-**H**; n=11) embryos. Slices from confocal Z-stack. (**I,J**) Profile plots of *etv5b* and *neurod4* HCR signal across the hindbrain of representative sibling control (*plzf*^*+/+*^;*plzfb*^*-/-*^) and *plzfa*^*-/-*^;*plzfb*^*-/-*^ embryos; anterior to the left. Scale bars: 50 µm.

Analysis of HCR signal profiles along the antero-posterior axis of the hindbrain confirmed that ectopic *etv5b* expression occurs outside rhombomere centres in the double homozygotes (Fig. 3I, J; Fig. S2). The expression pattern of *neurod4* was found to be altered in the *plzfa*^*-/-*^;*plzfb*^*-/-*^ hindbrain, typically with a loss of the characteristic ‘double peak’ flanking each rhombomere boundary (Fig. 3I, J; Fig. S2). These results show that the spatial pattern of neurogenesis is disrupted in embryos lacking Plzf. Since Fgf inhibits neurogenesis in rhombomere centres (Gonzalez-Quevedo et al., 2010), the altered pattern of neurogenesis upon loss of Plzf may be a consequence of excess and more widespread Fgf signalling.

### Loss of Plzf leads to excess Fgf signalling in the hindbrain at 24 hpf

From approximately 20 hpf, Fgf20 signalling from clusters of early-born neurons inhibits neurogenesis in rhombomere centres, resulting in neurogenic and non-neurogenic zones in the hindbrain (Gonzalez-Quevedo et al. 2010). Our finding that the Fgf target *etv5b* is ectopically expressed in the hindbrain at 30 hpf in embryos lacking Plzf prompted us to examine the relationship between Plzf and Fgf signalling at an earlier stage.

We examined zebrafish embryos with mutations in *plzfa* alone (*plzfa*^*-/-*^) and both *plzfa* and *plzfb* (*plzfa*^*-/-*^;*plzfb*^*-/-*^; see Materials and Methods) by staining for *etv5b* expression. In sibling controls, *etv5b* mRNA is confined to rhombomere centres at 24 hpf, whereas in *plzfa*^*-/-*^;*plzfb*^*-/-*^ double mutant embryos, *etv5b* expression is detected outside rhombomere centres (Fig. 4A, B; I, J). Ectopic *etv5b* expression is also observed in the anterior spinal cord, while the area of the hindbrain anterior to r3 appears relatively unaffected. By contrast, the *plzfa*^*-/-*^ single mutant shows a comparatively mild phenotype, with ectopic *etv5b* typically only detected in r4 (Fig. 4C, D). We therefore focused our subsequent analyses on the double mutant.

**Figure 4.**
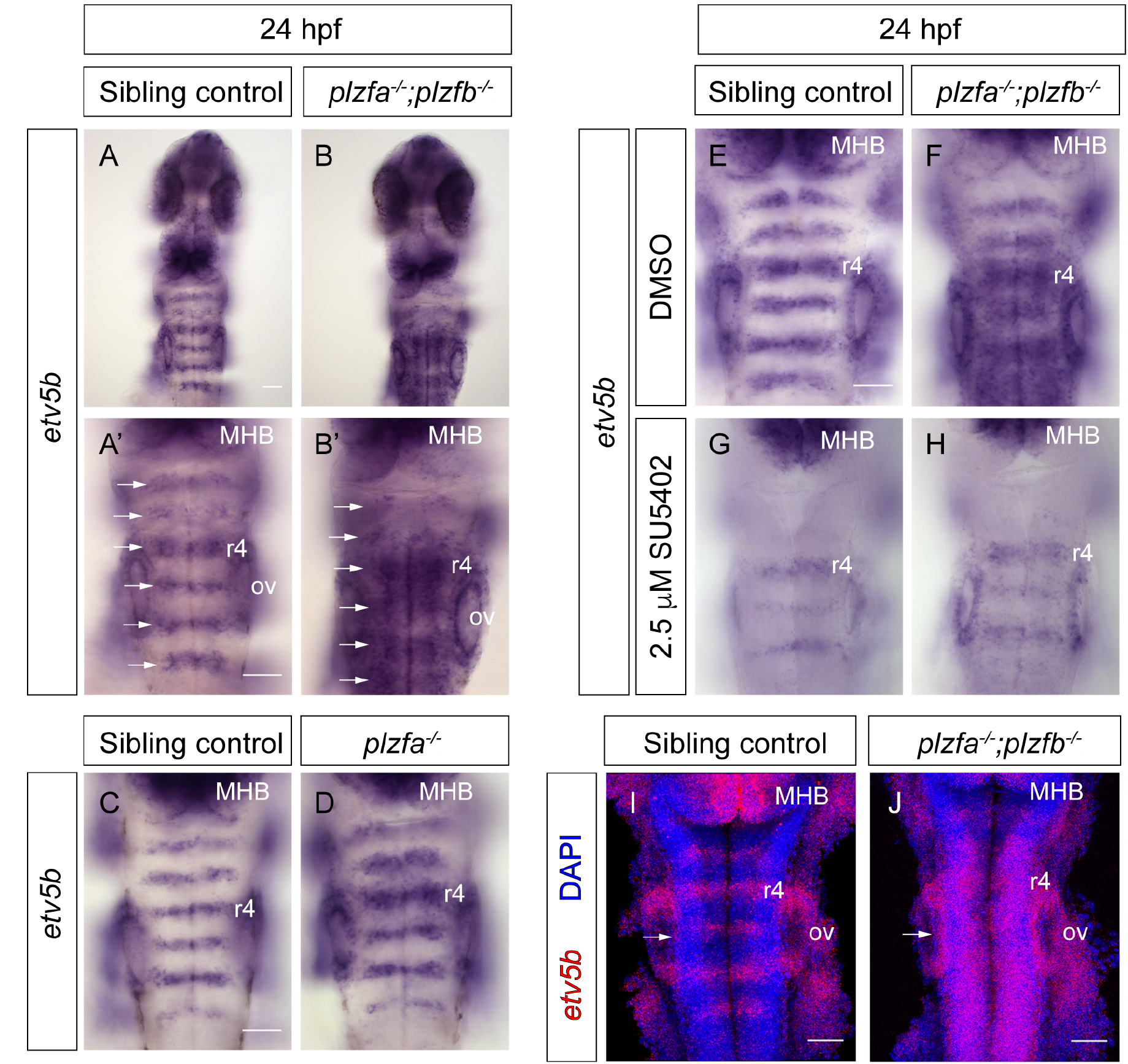
Spatial analysis of Fgf signalling in *plzf* mutants. (**A,B**) Colorimetric *in situ* hybridisation (ISH) for *etv5b* in sibling control (n=59) and *plzfa*^*-/-*^;*plzfb*^*-/-*^ (n=25) embryos, dorsal views of the hindbrain, wholemount. Arrows indicate rhombomere centres. (**C,D**) ISH for *etv5b* in sibling control (n=18) and *plzfa*^*-/-*^ (n=6) embryos. (**E,F**) Sibling control (E; n=88) and *plzfa*^*-/-*^;*plzfb*^*-/-*^ (F; n=28) embryos treated with DMSO from 22 to 24 hpf and stained for *etv5b*. (**G,H**) Sibling control (G; n=73) and *plzfa*^*-/-*^;*plzfb*^*-/-*^ (H; n=38) embryos treated with 2.5 µM SU5402 from 22 to 24 hpf and stained for *etv5b*. Three independent experiments. (**I,J**) HCR for *etv5b* in 24 hpf; sibling control (n=14) and *plzfa*^*-/-*^;*plzfb*^*-/-*^ (n=9) mutant embryos. 3D reconstructions of the Z-stack, dorsal view. Abbreviations: r, rhombomere; MHB, midbrain-hindbrain boundary; ov, otic vesicle. Scale bars: 50 µm

To determine whether the ectopic expression *of etv5b* observed after Plzf loss of function is a consequence of excess Fgf signalling, we inhibited Fgf receptor signalling in *plzfa*^*-/-*^;*plzfb*^*-/-*^ embryos with SU5402 (Mohammadi et al., 1997). Treating embryos from 22 to 24 hpf with 2.5. µM SU5402 was sufficient to abolish *etv5b* expression partially or completely in the hindbrain of both sibling controls and *plzfa*^*-/-*^;*plzfb*^*-/-*^ embryos, compared to vehicle controls (Fig. 4E-H). Together, these results demonstrate that loss of both *plzf* paralogues leads to excess Fgf signalling in the hindbrain at 24 hpf, suggesting a role for Plzf in negatively regulating Fgf signalling or expression.

### The expression of *fgf3* persists in the Plzf mutant hindbrain

A possible explanation for ectopic Fgf signalling in the Plzf double mutants is the ectopic expression of one or more Fgf ligands in the hindbrain. In the wild-type, at 24hpf, *fgf20a* is expressed in clusters of early-born neurons located bilaterally in the centre of each rhombomere, and accounts for the majority of *etv5b* expression in rhombomere centres (Gonzalez-Quevedo et al., 2010). Staining for *fgf20a* did not reveal any difference in expression between *plzfa*^*-/-*^;*plzfb*^*-/-*^ and control embryos. (Fig. 5A, B). The paralogous gene *fgf20b* is typically expressed in one or a few neurons within each *fgf20a*-expressing cluster, and this is also the case in *plzfa*^*-/-*^;*plzfb*^*-/-*^ mutants (Fig. 5C, D). Therefore, a broader source of Fgf20 signalling does not account for the phenotype observed in embryos lacking Plzf.

**Figure 5.**
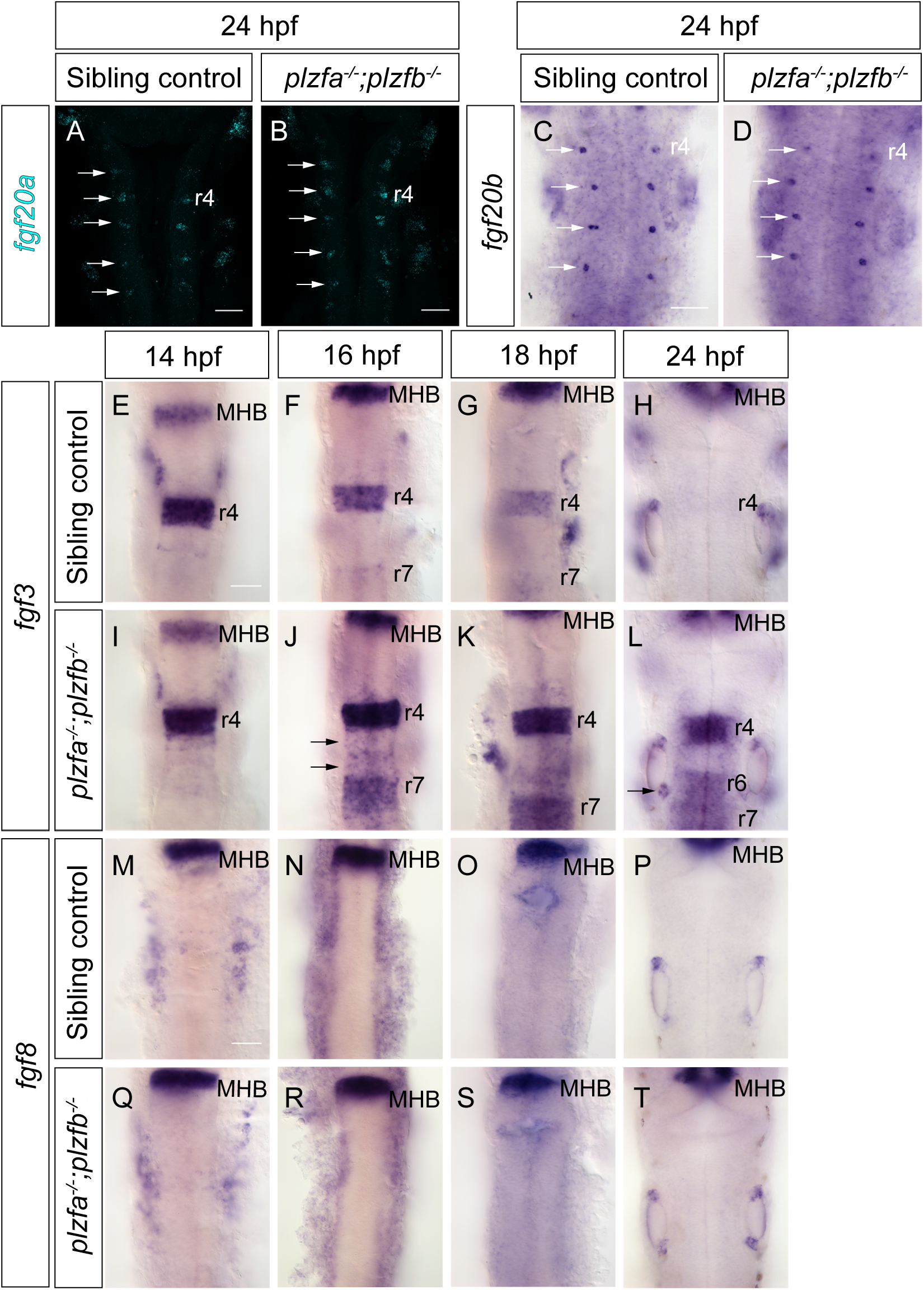
Expression of Fgf ligands in *plzf* mutants. (**A,B**) HCR for f*gf20a* in 24 hpf sibling control (n=26) and *plzfa*^*-/-*^;*plzfb*^*-/-*^ (n=12) embryos. 3D reconstructions of the Z-stack, dorsal view; arrows indicate *fgf20-*expressing neuronal clusters in rhombomere centres. (**C,D**) Colorimetric ISH for *fgf20b* in 24 hpf sibling control (n=15) and *plzfa*^*-/-*^;*plzfb*^*-/-*^ (n=4) embryos. Arrows indicate *fgf20-*expressing neuronal clusters in rhombomere centres. (**E**-**L**) ISH time-course of *fgf3* expression in the sibling control (E-H; 14 hpf: n=19; 16 hpf: n=26; 18 hpf: n=53; 24 hpf: n=47) and *plzfa*^*-/-*^;*plzfb*^*-/-*^ (I-L; 14 hpf: n=6; 16 hpf: n=10; 18 hpf: n=20; 24 hpf: n=12) hindbrain. Arrows in (J) indicate ectopic expression in r5/r6, arrow in (L) indicates ectopic expression in the posterior otic vesicle. (**M**-**T**) ISH time-course of *fgf8* expression in the sibling control (O-R; 14 hpf: n=39; 16 hpf: n=13; 18 hpf: n=16; 24 hpf: n=35) and *plzfa*^*-/-*^;*plzfb*^*-/-*^ (S-V; 14 hpf: n=8; 16 hpf: n=8; 18 hpf: n=5; 24 hpf: n=10) hindbrain. Abbreviations: r, rhombomere; MHB, midbrain-hindbrain boundary. Scale bars: 50 µm

An alternative source of ectopic Fgf signalling could be a ligand which is normally transiently expressed at an earlier stage of development and which persists in the mutant. In zebrafish, *fgf8* and *fgf3* are expressed in the prospective hindbrain from late gastrulation in a domain that becomes restricted to r4. These Fgf family members are required for correct segmental gene expression in the adjacent rhombomeres, as well as induction of the inner ear (Phillips et al., 2001; Maroon et al., 2002; Maves et al., 2002; Walshe et al., 2002). *fgf8* is downregulated in r4 soon after hindbrain segmentation (Walshe et al., 2002), whereas *fgf3* expression persists until later and is downregulated by approximately 19 hpf (Maroon et al., 2002; Reuter et al., 2019).

To determine whether *fgf3* or *fgf8* expression persists in the hindbrain, we examined the expression of each gene in *plzfa*^*-/-*^;*plzfb*^*-/-*^ mutant embryos and sibling controls at 14 hpf, 16 hpf, 18 hpf and 24 hpf. At 14 hpf, *fgf3* is expressed in r4 at comparable levels in both mutants and controls (Fig. 5E, I). At 16 hpf, in contrast, *plzfa*^*-/-*^;*plzfb*^*-/-*^ embryos show high levels of *fgf3* in r4 compared to controls, and ectopic *fgf3* expression is observed in r5 and r6 (Fig. 5F, J). In sibling controls, low levels of *fgf3* are consistently detected in r7 and the anterior spinal cord at 16hpf while expression levels in these domains are higher in the double mutants, (Fig. 5F, J). At 18 hpf, ectopic *fgf3* persists in r5 and r6 and *fgf3* levels remain high in r7 and the anterior spinal cord (Fig. 5G, K). At this stage, the sibling controls are downregulating *fgf3* in r4, whereas high levels persist in r4 in the double mutants (Fig. 5G, K.) By 24 hpf, no *fgf3* expression can be detected in the sibling controls, but expression persists in *plzfa*^*-/-*^;*plzfb*^*-/-*^ in r4, at intermediate levels in r6, r7 and the anterior spinal cord, and at low levels in r5 (Fig. 5H, L). *fgf3* expression in r7 and the anterior spinal cord has not been described previously in wild-type embryos; since the sibling controls lack functional *plzfb*, the low levels of *fgf3* detected at these axial levels could be a consequence of *plzfb* loss of function. We therefore examined *fgf3* expression in wild-type embryos at 16 hpf and found that low levels are present in r7 and the anterior spinal cord (Fig. S3), suggesting that the ectopic *fgf3* detected in these tissues in the Plzf double mutant at 24 hpf represents a failure to downregulate *fgf3* in its normal expression domain. By contrast, *fgf8* expression is not detected in the hindbrain of sibling controls or double mutants at any of the stages examined (Fig. 5O-V).

Taken together, these results indicate that Plzf downregulates *fgf3* expression in the hindbrain between 14 hpf and 24 hpf. This timing is consistent with the expression pattern of Plzf, with the onset of higher levels of Plzf protein in the posterior hindbrain (Fig.1C, F, I) preceding the downregulation of *fgf3* (Fig. 5E-H). The expression pattern of *fgf3* in 24 hpf embryos lacking Plzf is consistent with Fgf signalling being less affected in the anterior hindbrain (Fig. 4). Although we cannot exclude the contribution of other Fgf ligands, our results suggest that ectopic *fgf3* is responsible for the ectopic Fgf signalling observed in the absence of Plzf.

### The inner ear is transiently anteriorised in embryos lacking Plzf

The inner ear develops adjacent to rhombomeres 4-6 and is dependent on the hindbrain for several aspects of its development. In zebrafish, induction of the inner ear requires Fgf3 and Fgf8 signalling from r4 among other sources (Phillips et al., 2001; Maroon et al., 2002); later, Fgfs from the hindbrain act as an anteriorising signal to pattern the otic vesicle (Kwak et al., 2002; Hammond and Whitfield, 2011; Hartwell et al., 2019). In addition to the ectopic *etv5b* expression detected in the hindbrain of *plzfa*^*-/-*^;*plzfb*^*-/-*^ embryos, we also observe ectopic *etv5b* in the medial wall of the otic vesicle at 24 hpf (Fig. 6A, B). Thus, the otic vesicle may receive an excess of Fgf which in turn may result in its anteriorization. We therefore assayed sibling control and Plzf double mutant otic vesicles for the expression of the anterior markers *pax5* and *hmx2. pax5* is expressed in the utricular macula (Kwak et al., 2006) and expands posteriorly in response to Fgf misexpression (Kwak et al., 2002; Hammond et al., 2011). Indeed, we find an enlarged domain of both *pax5* and *hmx2* expression in *plzfa*^*-/-*^;*plzfb*^*-/-*^ embryos, with expression detected throughout the medial wall of the otic vesicle (Fig. 6C, D; I, J). In contrast, ectopic expression of *pax5* is never detected in the *plzfa*^*-/-*^ otic vesicle (Fig. 6G, H). Since *plzfa*, but not *plzfb* is expressed in the otic vesicle (Fig. 1), this result argues against a tissue-autonomous phenotype in the inner ear, but instead suggests that the expansion of anterior markers is due to increased signalling from the hindbrain. Like *pax5* and *hmx2*, the ventro-lateral otic vesicle marker *otx1* (Li et al., 1994) depends on Fgf signalling and is upregulated in response to ubiquitous misexpression of Fgf3 (Maier and Whitfield, 2014). However, no difference in *otx1* expression was detected between sibling controls and embryos lacking both *plzf* paralogues (Fig. 6K, L). This indicates that the patterning defects observed at 24 hpf are specific to the medial wall of the inner ear and further supports the conclusion that excess Fgf from the hindbrain anteriorises the inner ear in Plzf mutants.

**Figure 6.**
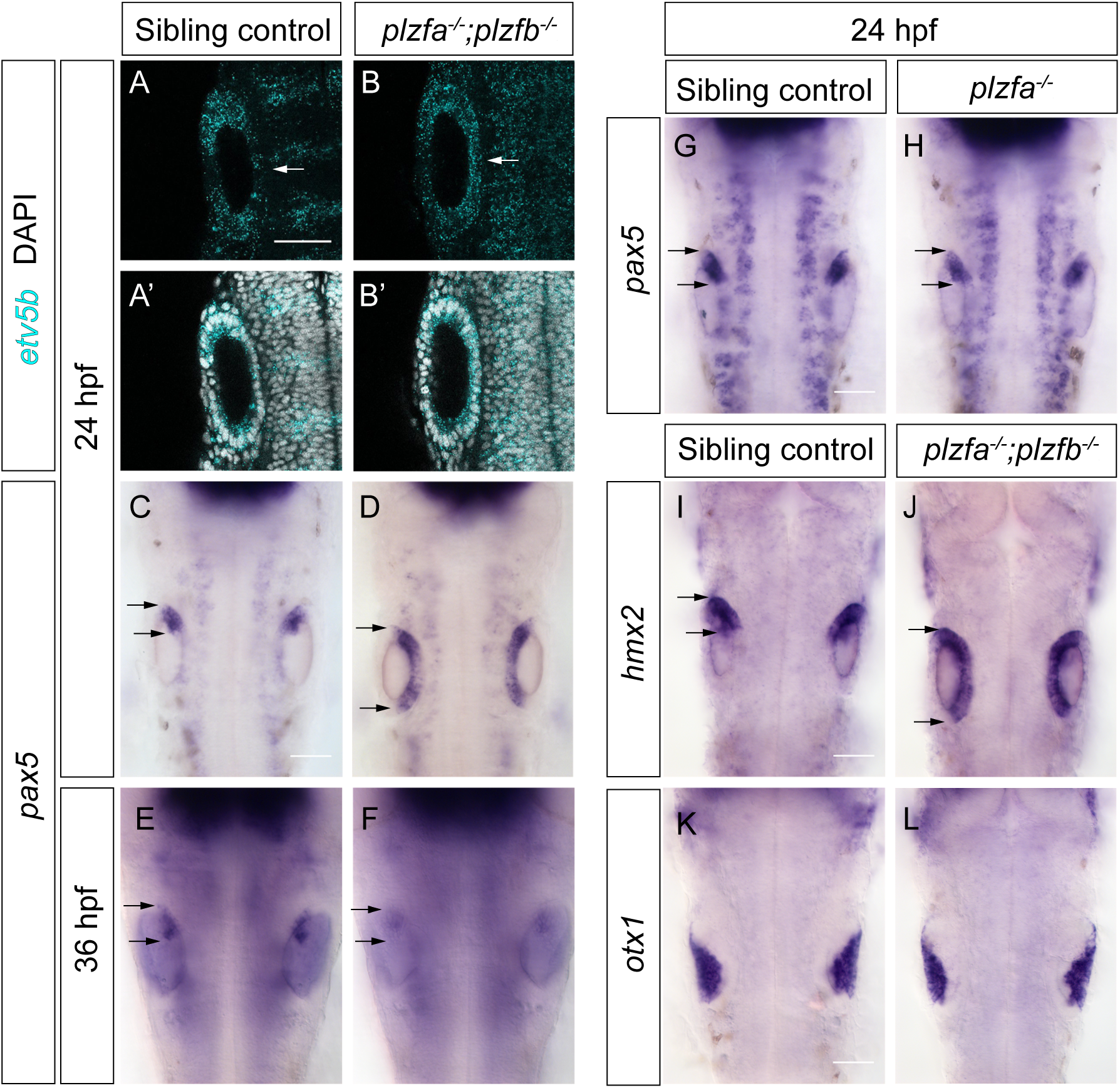
Inner ear patterning in *plzf* mutants. (**A,B**) HCR for *etv5b* in 24 hpf sibling control (n=14) and *plzfa*^*-/-*^;*plzfb*^*-/-*^ (n=9) embryos. Confocal Z-stack slices from experiment shown in Fig. 2 I,J. Arrows indicate the medial wall of the otic vesicle. (**C,D**) Colorimetric ISH for *pax5* in 24 hpf sibling control (n=10) and *plzfa*^*-/-*^;*plzfb*^*-/-*^ (n=11) embryos. Arrows in (C-J) show the antero-posterior extent of marker gene expression in the otic vesicle. (**E,F**) ISH for *pax5* in 36 hpf sibling control (n=29) and *plzfa*^*-/-*^;*plzfb*^*-/-*^ (n=11) embryos. (**G,H**) ISH for *pax5* in 24 hpf sibling control (n=15) and *plzfa*^*-/-*^ (n=9) embryos. (**I,J**) ISH for *hmx2* in 24 hpf sibling control (n=40) and *plzfa*^*-/-*^;*plzfb*^*-/-*^ (n=10) embryos. (**K,L**) ISH for *otx1* in 24 hpf sibling control (n=25) and *plzfa*^*-/-*^;*plzfb*^*-/-*^ (n=6) embryos. Scale bars: 50 µm

Overexpression of Fgf3 between 14 and 18 hpf generates full mirror image anterior duplications of the otic vesicle, with *pax5* expression resolving into distinct anterior and posterior domains by 36 hpf (Hartwell et al., 2019). To determine whether the anteriorisation of the otic vesicle observed in Plzf mutants at 24 hpf results in an anterior duplication, we stained *plzfa*^*-/-*^;*plzfb*^*-/-*^ embryos for *pax5* mRNA at 36 hpf. Unlike at 24 hpf, *pax5* expression was confined to the anterior macula in both sibling control and *plzfa*^*-/-*^;*plzfb*^*-/-*^ otic vesicles, although its levels were consistently lower in the double mutant (Fig. 6E, F). Finally, no gross morphological defects of the inner ear were observed (data not shown). Together, these results suggest that the inner ear is transiently anteriorised in the absence of Plzf. This anteriorisation is likely the consequence of ectopic Fgf signalling in the hindbrain, consistent with the known role for Fgf3 in axial patterning of the otic vesicle, but does not lead to persistent patterning defects.

## Discussion

Embryonic tissues are sequentially patterned by signals which have stage-specific roles and may act through shared downstream pathways, necessitating control of where and when the expression of signalling pathway components is turned on and off during development. Here, we show that the transcription factor Plzf is required for switching between stage-specific patterns of Fgf signalling in the zebrafish hindbrain. Plzf protein expression is anticorrelated with *fgf3* expression, and in the absence of Plzf, *fgf3* expression fails to be downregulated, interfering with the patterning of neurogenesis and transiently disrupting antero-posterior patterning of the inner ear. These results also point to a difference in how the pattern of Fgf signalling in the hindbrain is regulated in amniotes and teleosts.

### Regulation of *fgf3* expression by Plzf

The expression of Fgf ligands in the zebrafish hindbrain is dynamic and Fgf signalling has different functions at different developmental stages. Our results suggest that the onset of higher levels of Plzf expression in the posterior hindbrain and anterior spinal cord leads to downregulation of *fgf3*. However, the mechanism by which Plzf represses *fgf3* is unknown. The homeobox transcription factor *hoxb1* is a known regulator of r4 identity (Studer et al., 1996; McClintock et al., 2002) and positively regulates *fgf3* expression in r4 (Waskiewicz et al., 2002; Choe et al., 2011; Weicksel et al., 2014). GATA2 and GATA3 specify r4 identity downstream of Hoxb1 in the mouse hindbrain (Pata et al., 1999), and GATA4 mediates *fgf3* upregulation upon retinoic acid-induced differentiation of F9 embryonal carcinoma cells into parietal endoderm (Murakami et al., 1999), a cell type which also expresses *hoxb1* (Boylan et al., 1993). In a screen for downstream effectors of zebrafish *hoxb1b*, the protein phosphatase 1 regulatory subunit *ppp1r14al*, a potential regulator of GATA activity, was found to positively regulate *fgf3* expression in r4 (Choe et al., 2011) Intriguingly, Plzf binds the mouse GATA4 regulatory region and upregulates its expression in heart cells (Wang et al., 2012b). GATA factors are thus candidates for Plzf targets in r4. So far, there is no experimental evidence for direct Plzf regulatory targets in the hindbrain or elsewhere in the CNS. Studies that assay Plzf binding genome-wide in neural tissue will likely shed light on the mechanism of *fgf3* downregulation by Plzf.

### Regulation of Plzf expression

The sequential onset of Plzf expression first in the anterior CNS, and then more posteriorly, suggests that regulation of Plzf levels is coupled to an antero-posterior patterning process. During early hindbrain development, retinoic acid (RA) produced in the somites acts as a posteriorizing signal and is thought to form a gradient across the hindbrain (reviewed in Schilling et al., 2016; Frank and Sela-Donenfeld 2019).

The hindbrain RA gradient has been proposed to be generated by rhombomere-specific expression of RA-metabolising Cyp26 enzymes (Sirbu et al., 2005; Hernandez et al., 2007; White et al., 2007). In one model, the dynamic expression of different *cyp26* genes in the zebrafish hindbrain leads to a progressive posterior shift of the anterior border of the RA-responsive domain during somitogenesis stages (Hernandez et al., 2007). This in turn suggests a model where low levels of RA signalling are permissive to Plzf expression, which is upregulated in the posterior hindbrain as the domain of active RA signalling recedes. Other signalling pathways that regulate hindbrain antero-posterior patterning, such as Fgf and Wnt, as well as downstream transcription factor networks (reviewed in Parker and Krumlauf, 2017; Frank and Sela-Donenfeld 2019) may also act upstream of Plzf.

### Plzf and the Fgf signalling switch in fish and amniotes

The hindbrain expression pattern of Plzf in zebrafish differs from that of amniotes. Both zebrafish and chicken have segmental expression of Plzf during early hindbrain development. However, in zebrafish, r1 and r5-6 have higher levels of Plzf expression, whereas the even-numbered rhombomeres are enriched for *PLZF* mRNA in chicken and mouse embryos (Cook et al., 1995). At later stages, *PLZF* is expressed in the rhombomere boundaries in chicken, mouse and rat (Cook et al., 1995; Takahashi and Osumi, 2011). In chicken embryos, the boundary cells constitute populations of slowly dividing, multipotent neuronal progenitors (Peretz et al., 2016). Here, we show that from approximately 24 hpf, Plzf is expressed in the ventricular zone of the zebrafish hindbrain and is absent from most post-mitotic neurons. Therefore, in both fish and amniotes, Plzf expression is initially rhombomere-specific and confined to neuronal progenitors at later stages.

We found that *fgf3* expression in the zebrafish hindbrain is anticorrelated with high levels of Plzf, and *fgf3* expression persists in the absence of Plzf. Notably, in amniotes, the expression pattern of Plzf correlates with that of *fgf3*, the latter being enriched initially in even-numbered rhombomeres and then gradually confined to rhombomere boundaries (Mahmood et al., 1995; Weisinger et al., 2008, 2010, 2011; Sela-Donenfeld et al., 2009). This suggests that the repression of *fgf3* by Plzf is not a feature conserved between amniotes and fish. In amniotes, two different mechanisms have been implicated in downregulating *fgf3* expression in rhombomere centres: Weisinger et al. (2008) showed that BMP signalling can repress *fgf3* and that local inhibition of BMP signalling by follistatins is required to maintain *fgf3* expression at rhombomere boundaries, while Sela-Donenfeld et al. (2009) demonstrated a requirement for a secreted factor diffusing from the boundaries themselves in downregulation of *fgf3* in non-boundary regions. These studies point to differential regulation of *fgf3* in boundaries versus the rest of the rhombomeres. Thus, amniotes appear to have boundary-specific mechanisms for maintaining *fgf3* transcription, whereas zebrafish downregulate *fgf3* throughout the hindbrain by a mechanism involving Plzf.

During hindbrain development, a switch from an early pattern of Fgf signalling regulating rhombomere-specific properties to a later pattern regulating neurogenesis takes place. The early phase involves Fgf3 in both amniotes and fish (Weisinger et al., 2008, 2010; Maves et al., 2002; Walshe et al., 2002), whereas different Fgf ligands are utilized at the late phase: Fgf3 in amniotes (Weisinger et al., 2011) and Fgf20 in fish (Gonzalez-Quevedo et al., 2010). The use of different ligands may explain the distinct mechanisms for *fgf3* downregulation in the two groups. Precise patterning of neurogenic zones by Fgf20 requires downregulation of Fgf3 in the entire zebrafish hindbrain; upon loss of Plzf, *fgf3* persists and the ligand likely diffuses to the adjacent rhombomeres, as evidenced by ectopic *etv5b* in r3 of *plzfa*^*-/-*^ *;plzfb*^*-/-*^ embryos. Notably, homodimerization of Fgf20 and other ligands of the Fgf9 family increases their affinity to heparin, limiting their diffusivity (Kalinina et al., 2009) and the signalling range of Fgf9 in tissue is increased when dimerization is inhibited (Harada et al., 2009). The short-range activity of Fgf20 may therefore be necessary to achieve precise patterning of neurogenic zones within a rhombomere. It is currently not known why the spatial pattern of neurogenesis in the hindbrain differs between fish and amniotes, but this difference likely underlies the requirement for different Fgf ligands at neurogenic stages.

### The role of Plzf in neural progenitors

In human embryonic stem cell-derived neural stem cells as well as those expanded from human embryonic hindbrain, Plzf expression marks stages with a high capacity for progenitor maintenance and wide differentiation potential (Elkabetz et al., 2008; Tailor et al., 2013). Plzf-deficient mice have a smaller cerebral cortex with a reduced number of neurons, which may reflect a depletion of the neural stem cell pool (Lin et al., 2019). Consistent with these findings, we show that Plzf is expressed in the neural progenitors of the zebrafish hindbrain.

In addition to the ventricular zone, Plzf expression was detected in early differentiating progenitors migrating out of the ventricular zone. This is consistent with the model of Sobieszczuk et al. (2010), according to which, for neural differentiation to proceed, Plzf must be downregulated in progenitors selected to differentiate in the Notch-mediated lateral inhibition process. We found that Plzf protein and *plzfa* mRNA expression overlap with that of *neurog1* but are largely mutually exclusive with *neurod4*. At the onset of neuronal differentiation, the Plzf protein is targeted for degradation by the ubiquitin ligase adaptor protein Btbd6 (Sobieszczuk et al., 2010); in *Xenopus, XBTBD6* is a direct transcriptional target of NeuroD but not Neurogenin (Seo et al., 2007) and the knockdown of XBTBD6 affects the expression of late, but not early, neuronal markers (Bury et al., 2008). In the zebrafish hindbrain, *btbd6b* is expressed in differentiating progenitors outside the ventricular zone (data not shown). Together, these observations suggest a model where Plzf is expressed together with *neurog1* during the early stages of the neurogenic cascade and downregulated as cells migrate out of the ventricular zone and start expressing *neurod4*.

Plzf has been implicated in the maintenance of neural progenitors in the CNS through negative regulation of neurogenesis (Sobieszczuk et al., 2010; Gaber et al., 2013). One potential mechanism by which Plzf may antagonize neural differentiation is repression of proneural gene expression (Sobieszczuk et al., 2010), although this was not shown directly. In contrast, Gaber et al. (2013) demonstrate that Plzf opposes differentiation of neural progenitors in the chicken and mouse spinal cord by upregulating Fgf receptor 3 (FGFR3). Our results indicate that Plzf negatively regulates Fgf signalling in the hindbrain by repressing *fgf3*. In accordance with the results of Gonzalez-Quevedo et al. (2010), we find that high levels of Fgf signalling correlate with reduced neurogenesis in *plzfa*^*-/-*^*;plzfb*^*-/-*^ mutant fish. Interestingly, in both fish and amniotes, Plzf acts to regulate neuronal differentiation by regulating components of the Fgf signalling pathway – in the former case by downregulating the expression of an Fgf ligand, and in the latter by positively regulating Fgf receptor expression levels. However, we cannot rule out a contribution of cell-intrinsic regulation of neurogenesis by Plzf in neural progenitors; such a mechanism may be masked by the excess Fgf signalling. The possibility that Plzf acts directly on proneural gene expression may be addressed in the future by the analysis of transcriptional targets of Plzf in the hindbrain. The apparently paradoxical result that the loss of function of an antagonist of neurogenesis results in reduced neural differentiation may be explained by the earlier, stage-specific role of Plzf in downregulating an Fgf signal involved in segmental patterning of the hindbrain.

Therefore, our results reveal an important role for Plzf in ensuring that the pattern and levels of Fgf signalling in the hindbrain are appropriate for each developmental stage.

## Materials & Methods

### Zebrafish lines

Wild-type *Danio rerio*, Tg[*egr2b:H2B-Citrine*] (Addison et al., 2018), *plzfa* single mutant and *plzfa/plzfb* double mutant (see below) embryos were produced and staged according to hours post fertilisation and morphological criteria (Kimmel *et al*. 1995). *plzfa*^*-/-*^;*plzfb*^*-/-*^ (*plzfa*^*-/-*^) embryos were produced by in-crossing *plzfa*^*+/-*^;*plzfb*^*-/-*^ (*plzfa*^*+/-*^) adults and siblings were used as controls. The *plzfa* genotype of these embryos was confirmed post-staining by restriction fragment length polymorphism (RFLP) analysis of DNA isolated from individual embryos.

### Generation of the *plzfa/plzfb* mutant lines

Transcription activator-like effector nuclease (TALEN) constructs were designed to specifically target the *plzfa* and *plzfb* genes. Briefly, each TALEN array was designed according to the following criteria: 1) 16-20 base pair (bp) target sequence length; 2) a 14-17 bp spacer between the two arrays of the targeting pair; 3) presence of a thymine base immediately upstream of the target sequence; 4) absence of target sequence homology with other genes. A list of potential TALENs was generated using online software (https://boglab.plp.iastate.edu/; Cermak et al., 2011) and subsequent selection was performed manually.

TALEN construction was performed using the Golden Gate cloning technique designed for rapid generation of large constructs (Engler et al., 2009; Cermak et al., 2011). Plasmids were obtained from Addgene (Cat #1000000016) and TALENs were built using the 5-day protocol described by Cermak *et al*. (2011). Golden Gate-compatible destination vectors pCS2TAL3-DD and pCS2TAL3-RR (Dahlem et al., 2012) were obtained from Addgene (Cat. No. 37275 and 37276). Constructed plasmids were linearised with NotI restriction enzyme and capped mRNA was synthesised using the SP6 mMessage Machine Kit (Ambion). Equal amounts of left and right TALEN mRNA (100pg each) were injected into 1-cell stage zebrafish wild type zebrafish embryos.

The presence of TALEN-induced mutations was determined by High Resolution Melt (HRM) curve analysis of genomic DNA (gDNA) from 3 days post-fertilisation (dpf) embryos or fin clips from adult fish. Approximately 100 bp of gDNA was amplified around the target site. The primers used for amplification were 5’-GCCGTGTGGATTTCAGAGAC-3’ and 5’-GCGCATCTGATTAGCCTTGT-3’ for *plzfa*, and 5’-GTTCTGTGCGCATGAAACTC-3’ and 5’-CAGCCACCCTACAACTCTCC-3’ for *plzfb*. Triplicate 20 µl reactions containing 1 µl of gDNA solution were amplified and denatured in the presence of MeltDoctor HRM Dye (Applied Biosystems) using an Applied Biosystems 7900HT Fast Real-Time PCR System according to the manufacturer’s instructions. HRM data were analysed using the Applied Biosystems HRM Software v2.0.

To sequence individual alleles, approximately 500 bp of gDNA around the target site was amplified by PCR. The primers used for amplification were 5’-ACAAGAAAACGAACAACTGCAA-3’ and 5’-CTTGGAGCGTGGCAGTGTAG-3’ for *plzfa*, and 5’-CAGTTGCAGGAGCACTCAAG-3’ and 5’-AACCGCCATCTTGTATGGAA-3’ for *plzfb*. The PCR product was cloned into pGEM-T Easy (Promega), transformed into competent bacteria, and individual colonies were picked to carry out colony PCR using SP6 and T7 primers. The resulting PCR product was sequenced (GATC Biotech).

Embryos injected with TALENs targeting *plzfa* were grown to adulthood and outcrossed to wild type fish in order to produce F1 embryos heterozygous for *plzfa* mutations. HRM analysis confirmed the presence of indel mutations in both F1 embryos and adults. Mutant alleles from F1 fish were sequenced and fish with an 8-bp deletion approximately 50 bp downstream of the translation start site were used as founders for the *plzfa*^*+/-*^ line, which was maintained in a heterozygous state as *plzfa*^*-/-*^ fish do not reach adulthood.

Embryos injected with TALENs targeting *plzfb* were grown to adulthood and incrossed. HRM analysis confirmed the presence of indel mutations in F1 embryos. Heterozygous F1 adults were identified by RFLP analysis and mutant alleles were sequenced. Fish with identical 2-bp deletions at amino acid position 34 were incrossed to generate the *plzfb*^*-/-*^ line, which was maintained in a homozygous state.

Double mutant fish were generated by crossing *plzfa*^*+/-*^ fish to *plzfb*^*-/-*^ fish. The double mutant line was maintained as *plzfa*^*+/-*^;*plzfb*^*-/-*^.

### Immunofluorescence analysis of zebrafish embryos

Embryos were grown to the desired stage and fixed in 4% paraformaldehyde (PFA) in phosphate-buffered saline (PBS) at room temperature for 2-3 hours, dechorionated and washed several times in PBT (0.1% Triton X-100) before blocking for 1-2 hours at room temperature with 5% goat serum (GS) in PBT, 1% DMSO (10% GS for experiments described in Fig. 2). Primary antibodies were then added in 2% GS in PBT, 1% DMSO (5% GS for experiments described in Fig. 2) and embryos were incubated with gentle shaking overnight at 4µ C. Embryos were washed extensively with PBT at room temperature and incubated overnight at 4µ C with 2% GS in PBT, 1% DMSO (5% GS for experiments described in Fig. 2) containing the secondary antibodies. Embryos were counterstained with DAPI (1:1000) for 30 min at room temperature and washed extensively with PBT. For whole-mount imaging, embryos were mounted on a glass slide in 70% glycerol and coverslipped. For transverse sections, embryos were embedded in 4% agarose and sections with a thickness of 80–120μm were generated using a Leica VT1000 S Vibratome. Confocal Z-stacks were acquired with a Leica SP5, Leica SP8 (Fig. 1I) or Leica TCS SP2 (Fig. 2) confocal microscope.

A rabbit polyclonal antibody was raised against the purified BTB domain of the zebrafish Plzfa protein (Harlan Bioproducts). This antibody was subsequently found to recognise both Plzfa and Plzfb proteins (Fig. S1). The primary antibodies used for immunofluorescence in zebrafish embryos were: anti-Plzf (Rabbit IgG polyclonal; custom) at 1:500 (Fig.1) or 1:2000 (Fig. 2), anti-HuC/D (Mouse IgG_2b_; Molecular Probes A-21272) at 1:200 and anti-GFAP (Rabbit IgG polyclonal; Dako M076101-2). Secondary goat antibodies were Alexa Fluor conjugates (Invitrogen) used at 1:400 dilution.

### *In situ* hybridisation

Embryos were fixed in 4% PFA for 4 hours at room temperature, dehydrated in 100% methanol and stored at -20µ C. Embryos were rehydrated through a series of 75%/50%/25% methanol in PBS and *in situ* hybridisation (ISH) was carried out as described previously (Xu et al., 1994). Colour development was carried out using 5-bromo-4-chloro-3-indolyl-phosphate (BCIP) and 4-nitro blue tetrazolium chloride (NBT). Double mutant and sibling control embryos from incrosses of *plzfa*^*+/-*^ or *plzfa*^*+/-*^;*plzfb*^*-/-*^ fish were always processed together and the genotype of individual embryos was confirmed after staining. For whole-mount imaging, embryos were mounted on a glass slide in 70% glycerol and coverslipped. Images were acquired with a Zeiss Axioplan2 with an Axiocam HRc camera.

Two-colour fluorescent ISH was carried out using the Fast Blue/Fast Red detection system as previously described (Lauter et al., 2011). Detection of the fluorescent Fast Blue signal was carried by excitation with a 633 nm laser and detecting wavelengths greater than 650 nm. The Fast Red fluorescent signal was detected by excitation with a 561 nm laser and detecting wavelengths greater than 570 nm. Care was taken not to develop the Fast Blue signal for too long as the production of precipitate can obscure the subsequent development of the Fast Red signal.

The following probes were generated from cDNA clones by *in vitro* transcription: *neurog1, neurod4, etv5b* (Gonzalez-Quevedo et al., 2010); *fgf3* (Breau et al., 2012); *fgf20b* (gift of A. Nechiporuk); *otx1* (Li et al., 1994); *hmx2* (Feng and Xu, 2010). The template for the *plzfa* probe was generated by PCR using primers AGAAGATGACGAGGAGCGG and TTGCCACATAGCTCGCATC against the full-length cDNA clone (Sobieszczuk et al., 2010). The *fgf8* and *pax5* probe templates (based on Ma and Zhang, 2015) were generated from 28 hpf zebrafish cDNA by PCR. The reverse primers included a T7 promoter for direct *in vitro* transcription from the PCR product:

*fgf8* forward AATCCGGACCTACCAGCTTT
*fgf8* reverse GAAATTAATACGACTCACTATAGGCACATCCTGTGCTTCGCTTA
*pax5* forward TTGCGTCAGCAAGATACTGG
*pax5* reverse GAAATTAATACGACTCACTATAGGCGTGTGGCTGCGCTATAGTA

### Hybridisation Chain Reaction (HCR)

Embryos were raised to the desired stage, fixed in 4% PFA for 4 hours at room temperature, dehydrated in 100% methanol and stored at -20µ C. Embryos were rehydrated through a series of 75%/50%/25% methanol in PBS and HCR v3.0 was carried out following the standard protocol for zebrafish embryos (Choi et al. 2018). DAPI was added as a counterstain (1:1000 in 5xSSCT, 45 min at room temperature) and embryos were mounted in 70% glycerol for imaging. Confocal Z-stacks were acquired with a Leica SP5 confocal microscope (Leica SP8 Falcon for Fig. 1G-H). All probes, amplifiers and buffers were purchased from Molecular Instruments. Profile plots were generated in ImageJ from sum projections of individual Z-stacks. A line was drawn from the centre of rhombomere 2 to rhombomere 7 and the width was adjusted to the width of the hindbrain. Grey values were extracted using the Plot Profile tool. Plotting was performed in Prism 7. 3D reconstructions of Z-stacks were generated in Imaris 9.3.1.

### RNAscope

RNAscope^®^ ISH (Wang et al., 2012a) was carried out using the RNAscope Multiplex Fluorescent v2 system as described previously (Guglielmi et al., 2021; Economou et al. 2022). The *fgf3* probes (gift of C. Hill) were detected by incubating the embryos with Multiplex FL V2 HRP-C4 for 15 min at 40µ C, followed by incubation with tyramide (Sigma, #T2879) coupled to fluorescein-NHS ester (Thermo Scientific, #46410) for 25 min in the dark at room temperature. Embryos were counterstained with DAPI (1:1000 in PBS + 0.1% Tween) for 30 min at room temperature. Confocal Z-stacks were acquired with a Leica SP8 Falcon confocal microscope.

### Inhibitor treatments

Embryos from *plzfa*^*+/-*^;*plzfb*^*-/-*^ incrosses were grown to the 22 hpf stage, dechorionated manually, and transferred to a 6-well plate (approximately 30 embryos per well). 3 ml of embryo medium containing either 2.5 μM SU5402 (Sigma, SML0443) or the equivalent volume of DMSO was added to each well. The embryos were incubated for 2 hours at 28.5µ C and fixed immediately after incubation for analysis by ISH. Sibling controls and double homozygous embryos were processed together and genotyped individually post-staining.

### Cell culture, transfection and immunohistochemistry

HEK293 cells were cultured as described (Poliakov et al., 2008) and transfected with 1 µg of the appropriate plasmid using FuGENE HD transfection reagent (Promega) according to the manufacturer’s instructions. For immunofluorescence staining, cells were fixed 48h post-transfection in 4% PFA for 15 min at room temperature, washed with PBST, blocked for 1 hour in 4% goat serum and 2% bovine serum albumin, and stained with primary and secondary antibodies diluted in blocking buffer (Odyssey).

## Acknowledgements

We thank Justina Yeung, Monica Tambalo and Luca Guglielmi for advice on experimental techniques, Mohamed Ismail for purifying the Plzfa BTB domain peptide, Alex Nechiporuk for the *fgf20b* probe, Caroline Hill for the *fgf3* RNAscope probes, and the Crick Aquatics and Light Microscopy facilities for their excellent support. This work was supported by a studentship jointly funded by King’s College London and the Francis Crick Institute, which receives its core funding from Cancer Research UK (FC001217), the UK Medical Research Council (FC001217) and the Wellcome Trust (FC001217), and the BBSRC project grant BB/M006964/1 to AS.

---

**Figure S1.**
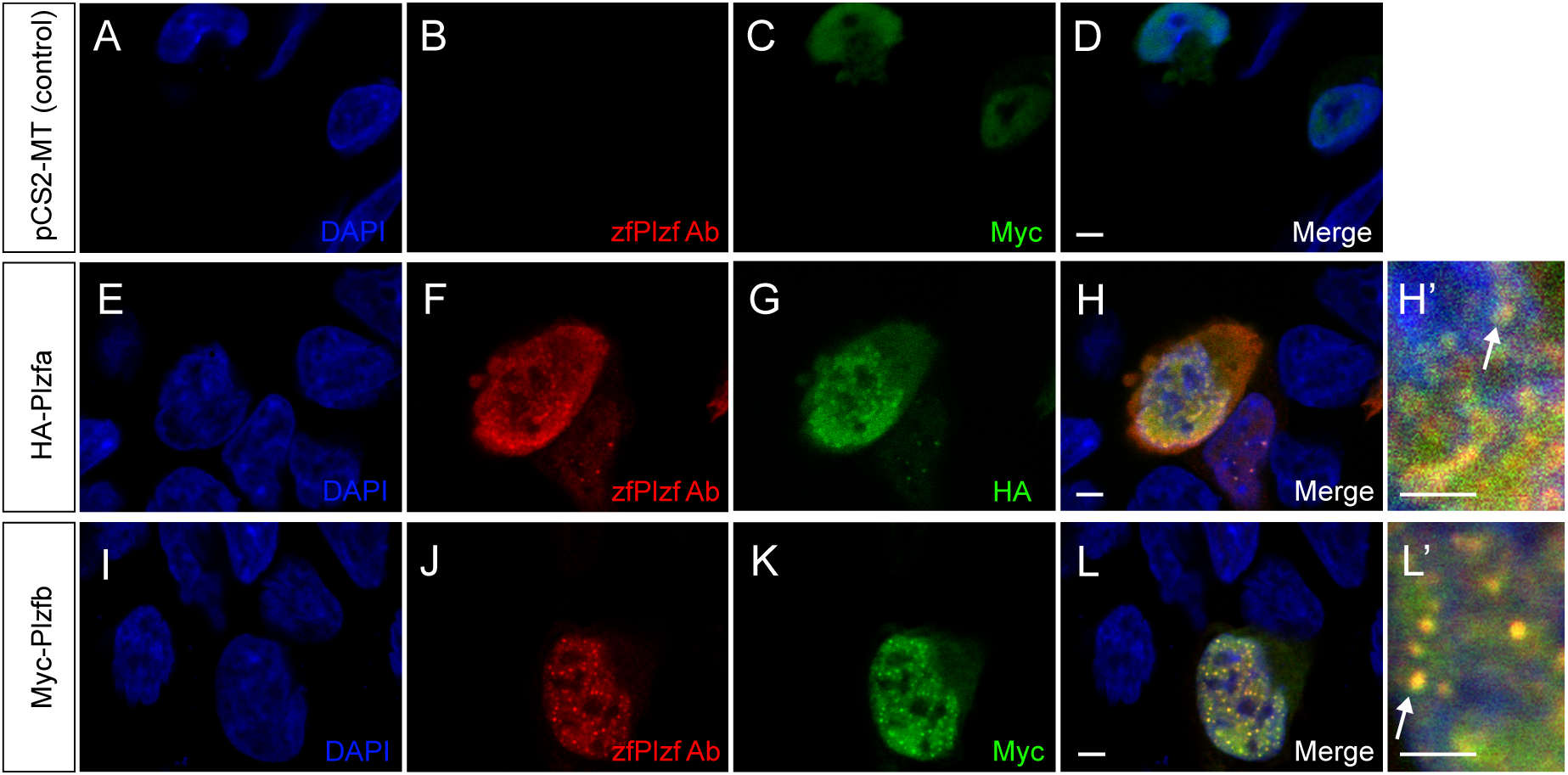
An antibody raised against the Plzfa BTB domain recognises both Plzf paralogues. Confocal images of HEK293 cells transfected with empty control vector (**A**-**D**), HA-tagged Plzfa (**E**-**H**) or Myc-tagged Plzfb (**I**-**L**) and stained with an antibody raised against the BTB domain of the zebrafish Plzfa protein (zfPlzf) and antibodies against the HA/Myc tags. Arrows point to nuclear specks showing colocalization of the zfPlzf and anti-HA/anti-Myc signals. Scale bars: 5 µm

**Figure S2.**
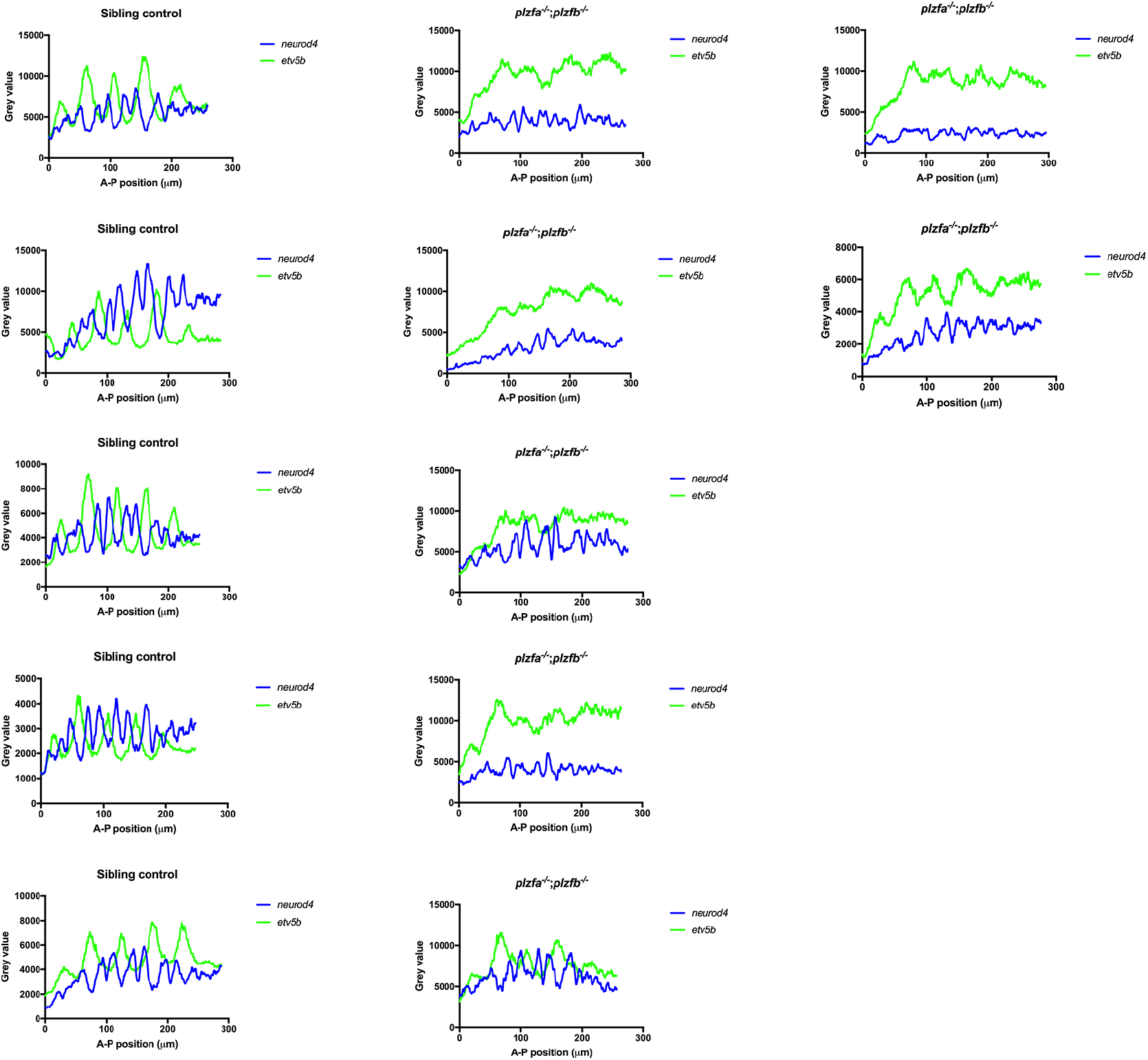
Pattern of neurogenesis in *plzf* mutants. Profile plots of *etv5b* and *neurod4* detected by HCR in individual 30h *plzfa*^*-/-*^;*plzfb*^*-/-*^ and sibling control *plzfa*^*+/+*^;*plzfb*^*-/-*^ embryos, as shown in Fig. 3. Anterior to the left.

**Figure S3.**
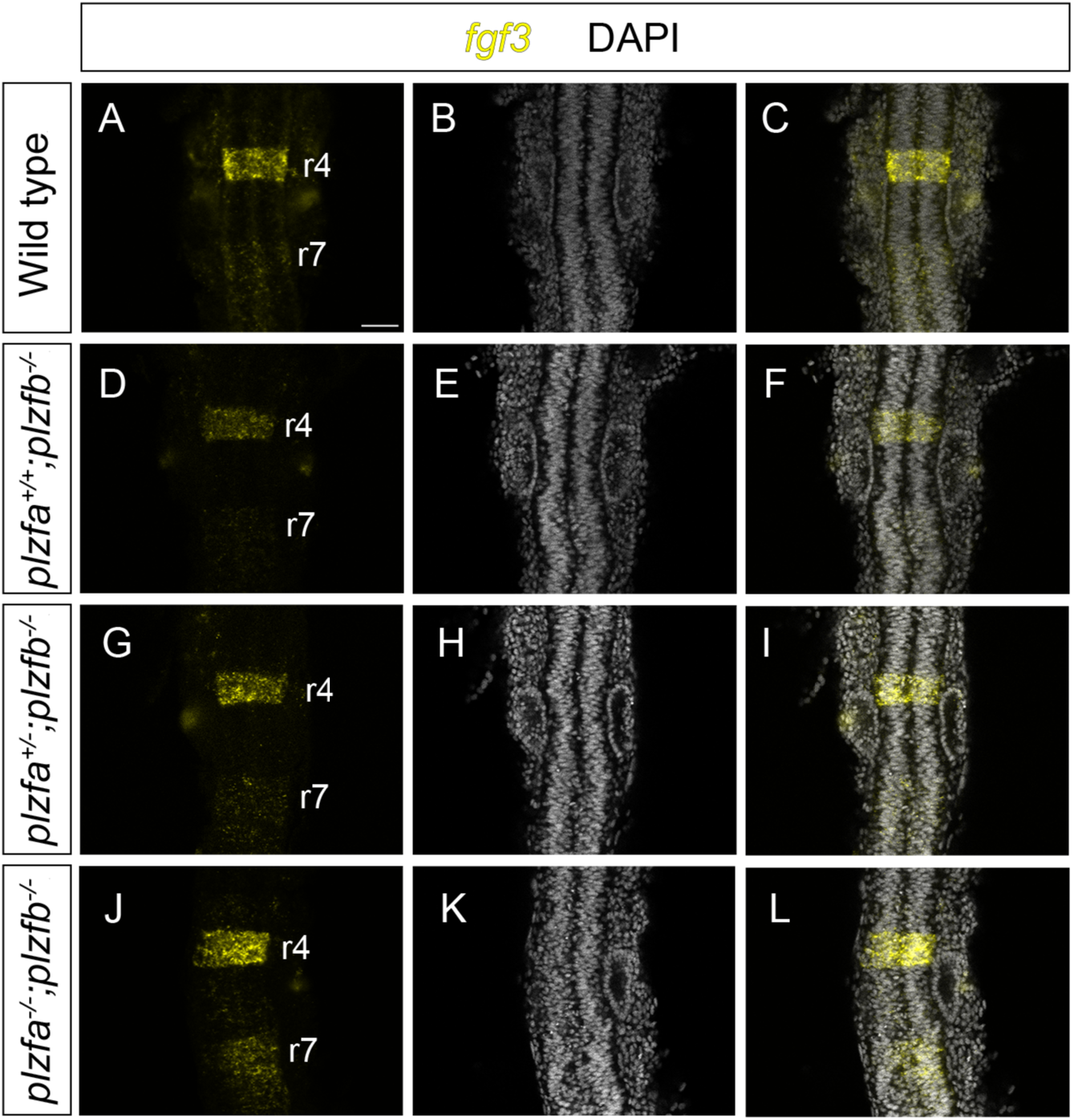
Expression of *fgf3* at 16 hpf. RNAscope staining against *fgf3* in wild type (**A**-**C**), *plzfa*^*+/+*^;*plzfb*^*-/-*^ (**D**-**F**), *plzfa*^*+/-*^;*plzfb*^*-/-*^ (**G**-**I**) and *plzfa*^*-/-*^;*plzfb*^*-/-*^ (**J**-**L**) embryos. Slices from confocal Z-stacks shown, anterior to the top. Abbreviations: r, rhombomere. Scale bar: 50 µm.

## Notes

### Competing Interest Statement

The authors have declared no competing interest.

